# Stretch-induced recruitment of myosin into transversal actin rings stabilizes axonal large cargo transport

**DOI:** 10.1101/2024.10.04.616720

**Authors:** Nizhum Rahman, Dietmar B. Oelz

**Affiliations:** School of Mathematics and Physics, The University of Queensland, QLD, 4072, Australia; Department of Mathematics, University of Michigan, Ann Arbor, MI 48109, United States

**Keywords:** actomyosin, axonal transport, stochastic simulation, tug-of-war, mechanotransduction

## Abstract

We study the axonal transport of large cargo vesicles and its feedback with contractile transversal actomyosin rings in axons through modelling and simulation. To this end, we simulate a mathematical model that integrates forces generated by the molecular motors and forces exerted by transversal actin rings. Our results predict that cargo vesicles exhibit bidirectional movement along with pauses in agreement with observations. It has been observed that during predominantly retrograde axonal cargo transport, blebbistatin treatment prolongs the periods spent by the cargo in anterograde transport. Our simulations show that this can be explained by mechanotransductive stretch-induced recruitment of myosin motors into transversal actin rings. These findings offer valuable insights into the complex dynamics of axonal cargo transport and propose potential avenues for further experimental research.

## 1 Introduction

Neuron cells are the backbone of information processing in the central nervous system. They exchange information and material through neurites, in particular through their long tubular axons. Axonal transport is the process by which motor proteins actively navigate microtubules (MTs) to transport a wide variety of cargoes such as vesicles and organelles from one end of the axon to the other [1]. These transport processes play a fundamental role for the development, function, and survival of nerve cells. Indeed, many diseases are associated with poor axonal transport such as Parkinson’s disease, Alzheimer’s disease, and Huntington’s disease [2–5].

Anterograde and retrograde transport refer to the movement of vesicles from the cell body towards the axon tip, and from the tip towards the cell body, respectively. Polar microtubule (MT) filaments serve as tracks [6]. Their minus ends tend to be found near the cell body, while MT plus ends are typically located in the periphery [7]. As a consequence, the direction of transport is determined by the motor proteins pulling the cargo. Members of the kinesin family are typically directed towards the plus end and therefore mostly involved in anterograde transport, whereas members of the dynein family are typically directed towards the MT minus ends and engaged in retrograde transport[7–11].

Axons may be as long as one meter compared to which vesicles move at a relatively slow speed [1]. The size of cargo vesicles (0.5 *−* 1.5µm [12]) may be significantly larger than the diameter of axons (e.g. 0.7µm [13] for connective axons in the corpus callosum [13]) in which case we refer to it as large cargo.

In the case of large cargo transport, transverse actomyosin rings immersed in the axonal cortex play a particular role. They are spaced at regular intervals, decorated with non-muscle myosin-II motor proteins and mechanically stabilize the axon [14]. In [15] (see Fig. 1A) it was reported that treatment with Blebbistatin (BLB) leads to

**Figure 1.**
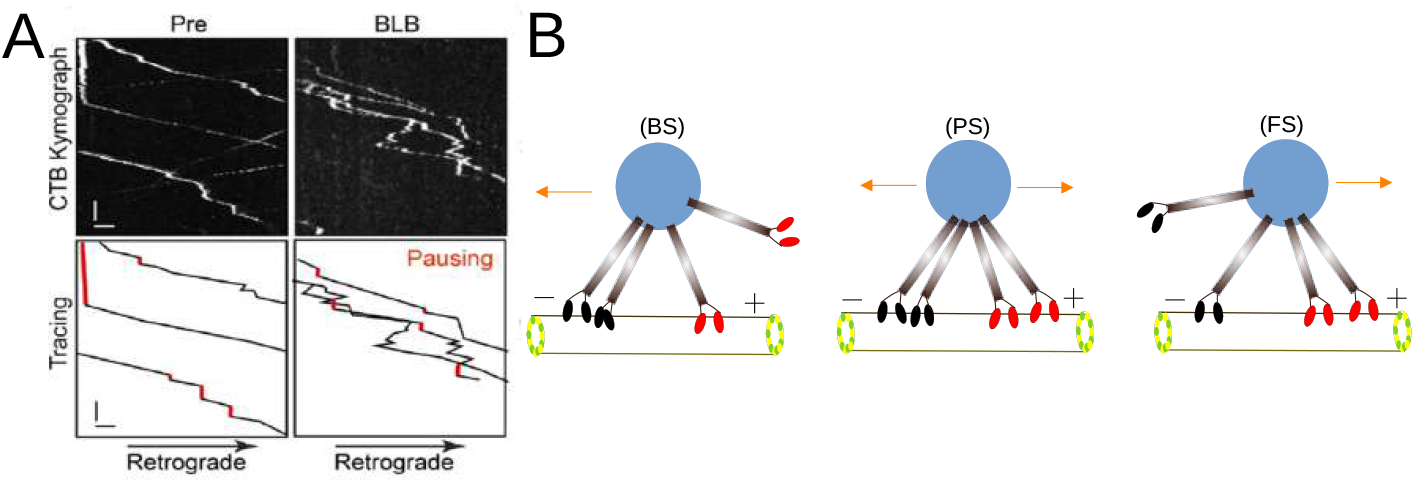
A: Kymographs of cargo vesicles in a single axon before (Pre) and after 60-min blebbistatin (+BLB) treatment. Tracings of retrograde cargoes are shown in the bottom panels. Pausing states with the trajectories are indicated with red. y-bar = 5 sec, x-bar = 1 µm (adapted from Figure 7d in [15]). B: Graphical sketch of stochastic model for bi-directional cargo transport. Cargo transported by two kinesins (red) and two dyneins (black) with the possible configuration (BS), (PS) and (FS). The configuration (BS) and (FS) represent the backward and forward motion respectively. The cargo movement is blocked if the numbers of antagonistic motors dynein and kinesin are equal.

1. an increase in the overall speed of cargo transport,
2. a decrease in the fraction of time spent in pause,
3. an increase in the fraction of time spent in anterograde transport during predominant retrograde transport and
4. a general increase of axon radius.

A novel mathematical model describing the interplay of actomyosin ring contraction, cargo propulsion and drag has been introduced in [16]. This model predicts an inverse relationship between cargo velocity and size. In agreement with experimental observations [15], it also predicts that large cargo moves at a slower speed in contrast to smaller cargo. It had, however, been formulated to explain the passive drag induced by the actomyosin rings on the cargo. For that reason, this model does not incorporate an accurate description of the motor protein forces driving cargo transport and of their stochastic nature.

In this study, we extend the model put forward in [16] coupling it with a classical model for the tug-of-war of molecular motor proteins[7, 17]. We carry out stochastic simulations to explore axonal cargo transport driven by randomly attaching and detaching motor proteins of the kinesin and dynein families. Our goal is to investigate the bidirectional movement of a large cargo due to molecular motors during axonal transport. It is our goal to explain the observations reported in [15] listed above. In particular, it is not clear how to explain that during predominantly retrograde axonal cargo transport, blebbistatin (BLB) treatment prolongs the periods spent by the cargo in anterograde transport. To address this question, we incorporate mechanotransduction through stretch-induced activation of myosin motors in transversal actin rings into our model and show that this is sufficient to explain these observations.

This study is structured as follows. We introduce the mathematical model for the tug-of-war of molecular motors engaged in transport of large cargo vesicles in section 2. In section 3 we detail the numerical scheme we use for our simulations. Stochastic simulations of tug-of-war during retrograde axonal transport are presented in section (4). Axonal transport relies not only on the motor proteins dynein and kinesins but it is crucial to recognize that actin filaments, in association with myosin, also contribute significantly to the transport of axonal cargo [18]. To further investigate the effect of BLB during the axonal cargo transport, we study the contractile feedback of actin ring stretching in the section (5).

## 2 Model for tug-of-war of antagonistic molecular motors

Our objective is to build on the mathematical model for axonal cargo transport, as established by Rahman and Oelz [16]. In this model. the axon is treated as a one-dimensional structure along which concentric rings with indices *j ∈*ℤ are positioned at equidistant locations *y*_*j*_ *∈* ℝ (see Figures 2 and 3).

**Figure 2.**
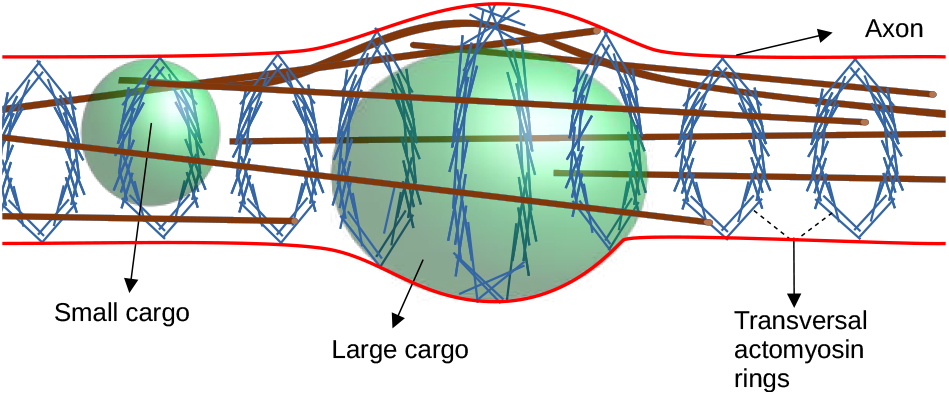
Actin rings (blue, sketched according to the closed ring hypothesis [14]) are positioned along the axon with equidistant spacing and wrapped around the circumference of the axon. Vesicles of various sizes (green) are moving through the axon. Microtubules (brown) provide transport rails for the molecular motor driven transport of vesicles. Adapted from [16].

**Figure 3.**
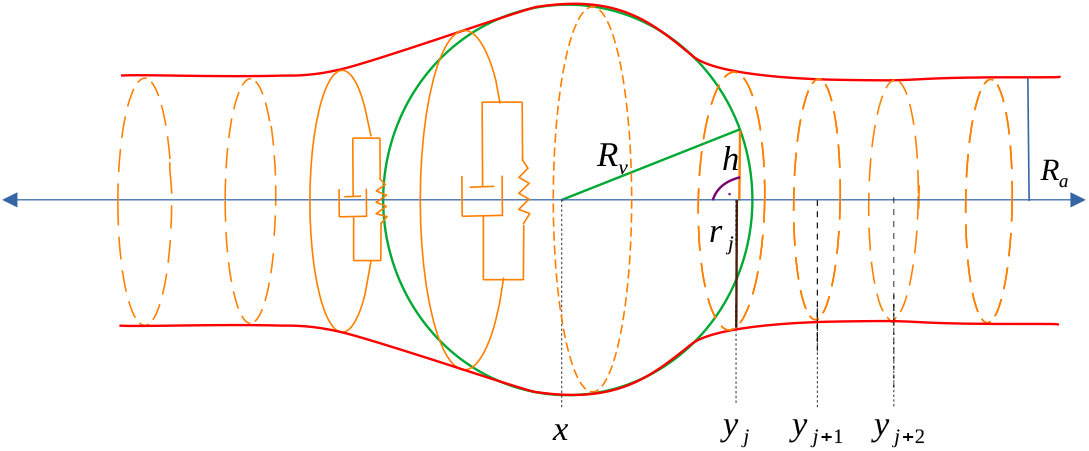
Sketch of the constituents of the mathematical model depicting a cargo vesicle at position *x ∈* ℝ (green with radius *R*_*v*_) moving from left to right through the axon with radius *R*_*a*_. As it is moving through the axon it extends the radii *r*_*j*_ of the actin rings uniformly positioned at *y*_*j*_, *y*_*j*+1_, etc. The function *h*_*j*_ denotes the “height” of the vesicle at the position of ring *j* (adapted from [16]).

The model’s time-dependent degrees of freedom are the position *x* = *x*(*t*) *∈* ℝ of a large cargo vesicle with radius *R*_*v*_ at time *t ≥* 0 and the radii of actin rings denoted by *r*_*j*_ = *r*_*j*_(*t*) *>* 0. Based on these quantities, the cross-sectional radius *h*_*j*_(*t*) of the vesicle at the position of the actin ring *j* is computed as a function of the distance between ring and vesicle position, *h*_*j*_(*t*) = *h*(*x*(*t*) *− y*_*j*_(*t*)) where

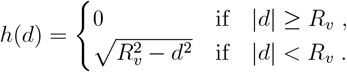

Since the vesicle is always contained in the axon, its cross-sectional radius at the position of a given actomyosin ring cannot be larger than the radius of that actin ring *j*. This introduces the constraint

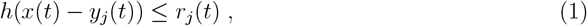

which is enforced by a force (Lagrange multiplier) in radial direction denoted by *λ*_*j*_(*t*).

The actin rings are treated as viscoelastic Kelvin-Voigt elements with equilibrium radius *R*_*a*_ (see Figure 3). The elastic force, illustrated in Figure 3 as a spring, is given by

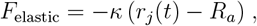

where *κ >* 0 represents the constant of elasticity. The resistance to changes in actin ring size, visualized in Figure 3 by a dashpot, is expressed by the viscous force

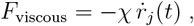

where *χ* represents the coefficient of viscosity and 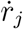 is the rate of change of the actomyosin ring radius. This coefficient characterizes the internal resistance of the actin rings to shortening and stretching, encompassing the stretching and realignment of actin fibers and cross-linker proteins, as well as transient unbinding from crosslinker proteins and myosin motors.

The resulting force-balance equation in radial direction for actomyosin ring *j* (see [16]) is given by

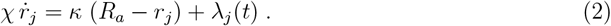

The force balance in the direction of the axon involves drag friction caused by the interaction of the vesicle with the cytoplasm and axonal cell membrane. It is modeled as linear drag with a positive coefficient *ξ >* 0,

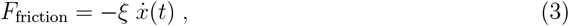

where 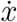 represents the velocity of the cargo vesicle.

The cargo vesicle is pulled by molecular motors pulling a cargo (see Fig 1B). The standard model for the force exerted by molecular motors is an affine force-velocity relation. Its parameters are the stall force and the free-moving velocity of the molecular motor [19, 20].

We assume that *N* motor proteins pull in anterograde direction [11] exerting the force

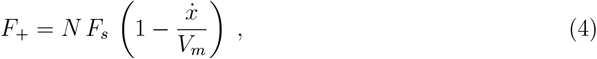

where *F*_*s*_ is the stall force of the molecular motor, *V*_*m*_ is its free moving velocity and 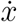 is the vesicle transport velocity. Similarly, the force *F*_*−*_ by which *M* motors are pulling the vesicle in retrograde direction is given by

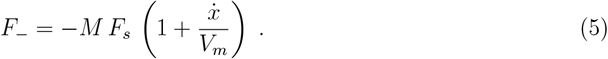

Combining both forces we get the motor force acting on the vesicle given by

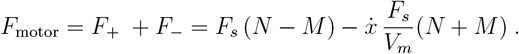

The force balance in the direction of the axon is then given by The system of equations is given by

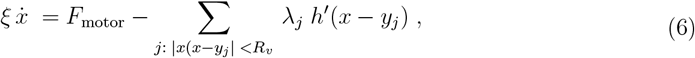

which involves the vertical component of the interaction force *λ*_*j*_ pressing upon the vesicle with slope *h*^*′*^(*x− y*_*j*_). The full model is the coupled system (2), (1), (6). For details of its derivation and its homogenisation limit for densily positioned actomyosin rings see [16].

For numerical computations, we make use of the fact that solutions can be approximated by the a variational time-stepping procedure where we compute the time-discrete solution *x*^*n*^, 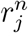 for every index of time *n* = 1, … with timesteps Δ*t* as a minimiser of an energy functional,

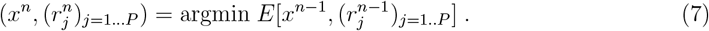

Here we assume the presence of *P* actomyosin rings and that the solution at the previous point in time 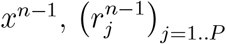 is known.

The energy functional involves conservative and dissipative energies, as well as a penalising potential for the constraint (1) (see [16]),

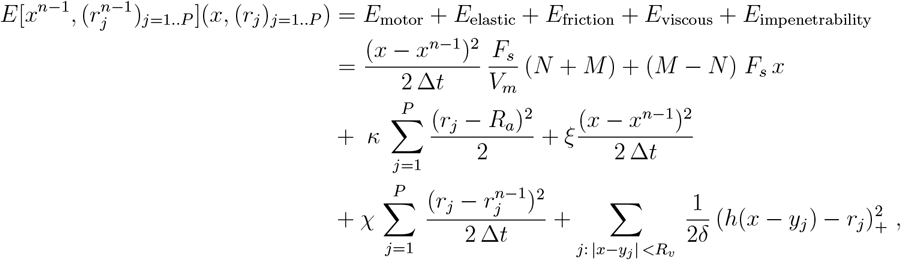

where 1*/δ* is an arbitrary large parameter for the penalising potential.

We also model the attachment of motor proteins to the MTs in retrograde and anterograde directions at a constant rate. We keep the setup simple by assuming that the force generation of molecular motors of both anterograde and retrograde direction is described by the same set of parameters. Yet, to mimic experimental observations reported in [15], we assume that the attachment rate for retrograde motors is higher than for anterograde motors (i.e.,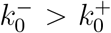), which implies that typically more motors will be pulling towards the cell body and that the vesicle tends to move in retrograde direction.

The motor proteins detach from the MTs at a rate which depends on the mechanical load. The detachment rate increases exponentially with the mechanical load acting on motors in anterograde direction (*F*_+_) and in retrograde direction (*F*_*−*_), respectively. The detachment rates are given by Bell’s law [7]

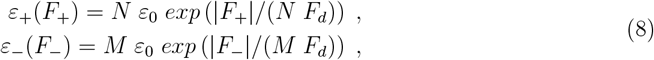

where *F*_*d*_ is the detachment force and *ε*_0_ the forceless detachment rate. Here, *ε*_+_ represents the motor detachment rate of motors pulling in anterograde direction whereas the motors pulling in retrograde direction detach with rate *ε*_*−*_.

During axonal transport, the number of motor proteins attached and detached from the MTs is changing randomly. As a consequence, at any time the cargo can move forward or backward or pause. Note that particularly in retrograde direction the on-rate is faster than the base off-rate *ϵ*_0_. For the entire population of attached motors, however, the off-rate scales linearly with their abundance (see Eq.(8)) and therefore the total number of motors remains bounded. For more details on how we simulate stochastic binding and unbinding of motors to microtubules (MT), see section 3.

Many of the parameter values listed in Table 1 may vary over various orders of magnitudes: For example, the diameter of axons in the central nervous system ranges from 0.1 *µm* to 10 *µm* [22]. The diameter of cargo that moves through the axon varies from 0.05 µm to 3 µm [15]. On the other hand, other parameters such as the spacing between the actin rings wrapped around the axon which is between 0.18 *µ*m to 0.19 *µ*m [21] hardly varies between different cell types.

**Table 1:** List of parameters values.

| Description | Symbol | Value | References/derivation |
| --- | --- | --- | --- |
| Radius of cargo vesicle | $R_v$ | 0.3 $\mu\text{m}$ | [15] |
| Equilibrium radius of axon | $R_a$ | 0.2 $\mu\text{m}$ | [13, 15] |
| The stall force | $F_s$ | 2.50 pN | [20] |
| Detachment force | $F_d$ | 2.0 pN | [20] |
| Attachment rate (anterograde direction) | $k_0^+$ | 1.0 $\text{sec}^{-1}$ | [20] |
| Attachment rate (retrograde direction) | $k_0^-$ | 3.0 $\text{sec}^{-1}$ | [20] |
| Forceless detachment rate | $\varepsilon_0$ | 1.0 $\text{sec}^{-1}$ | [20] |
| Free moving velocity | $V_m$ | 0.85 $\mu\text{m sec}^{-1}$ | [20] |
| Spacing between the actin rings | $\Delta y$ | 0.18 $\mu\text{m}$ | [21] |
| Drag friction | $\xi$ | 0.1 pN $\mu\text{m}^{-1} \text{sec}$ | see section 6 |
| Stretching stiffness of actin rings | $\kappa$ | 30 pN $\mu\text{m}^{-1}$ | see section 6 |
| Viscosity of actin rings | $\chi$ | 5 pN $\mu\text{m}^{-1} \text{sec}$ | see section 6 |

As for motor proteins, their characteristics depend on the type of motor proteins since both families of microtubule associated motor proteins – dynein and kinesin – have a large number of subtypes. Stall forces range from 1 pN to 6 pN [23], while detachment forces range from 0.87 pN to 4.01 pN [20]. The rate of attachment for motor proteins varies from 0.18*/s* to 5*/s*, and the rate of forceless detachment ranges from 0.26*/s* to 1.0*/s*

In axonal transport, the slower organelles move at a velocity of 0.1, *µm/s*, while the faster ones move at 1.0 *µm/s* [24]. [20]. In this study, however, we are interested in the regime that comes close to the experimental paper [15] in which axonal and vesicle radii are approximately given by the values in Table 1. For the parameters characterising motor proteins, we use average values so we do not have to distinguish between different types of molecular motors.

Finally, in this study we explicitly investigate the effect of variations on axonal stretching stiffness *κ*, axonal viscosity *χ*, drag friction *ξ* and axon radius. The values of these parameters in particular regimes are summarised in Table 2.

**Table 2:** The average values for axonal transport velocity and the fractions of time spent in anterograde motion, pause and retrograde motion.

| | Anterograde (%) | Pause (%) | Retrograde (%) | Velocity ($\mu\text{m sec}^{-1}$) |
| --- | --- | --- | --- | --- |
| reference | 9.24 | 17.70 | 73.06 | 0.35 |
| small $\kappa$ | 7.01 | 19.39 | 73.59 | 0.39 |
| large $\kappa$ | 13.35 | 28.04 | 58.61 | 0.24 |
| small $\chi$ | 8.92 | 8.79 | 82.29 | 0.51 |
| large $\chi$ | 9.47 | 26.28 | 64.24 | 0.22 |
| small axon | 11.35 | 21.98 | 66.67 | 0.23 |
| large axon | 8.09 | 12.47 | 79.44 | 0.49 |
| small $\xi$ | 9.05 | 18.74 | 72.21 | 0.36 |
| large $\xi$ | 9.93 | 20.29 | 69.78 | 0.30 |

An additional Table 2 has been provided to explore the effect of variations of the estimated parameter values listed in Table 1 on simulated axonal cargo transport.

## 3 Details of numerical scheme

We determine the length of timesteps Δ*t* and use the index *n* = 0, … to denote the numerical solution at time *n*Δ*t*. We start the simulation prescribing initial values for the vesicle position *x*^0^, the actomyosin ring radii 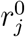 and the numbers of motors pulling the vesicle in anterograde direction given by *N* ^0^ as well as in retrograde direction given by *M* ^0^. In every timestep, we perform the following steps.

1. We find the position of the vesicle *x*^*n*^ and the radii of actomyosin rings 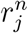 using numerical optimisation according to the variational scheme (7).
2. The forces *F*_+_ (in anterograde direction) and *F*_*−*_ (in retrograde direction) applied to the vesicle are obtained from the affine force-velocity relation for cargo transport [11] defined in Eq. 4 and Eq. 5 in which we approximate 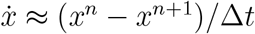.
3. We compute the motor detachment rates separately for motors pulling in anterograde and in retrograde direction. The detachment rates increase exponentially with the applied force [7] according to Eq. 8. Note that the attachment rates are assumed to be constant (see Table 1).
4. For the period of time Δ*t* we compute the number of motor proteins attaching and detaching to the cargo as follows:
  - The numbers of motor proteins attaching in either anterograde or retrograde direction (*M*) are sampled from the Poisson distribution with rates of 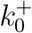 and 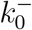, respectively.
  - The number of detaching motor proteins in anterograde and retrograde directions are sampled from the Poisson distribution, with detachment rates *ε*(*F*_+_) and *ε*(*F*_*−*_), respectively.
5. Repeat steps 1 *−* 3 for every time step Δ*t* until reaching *t*_*max*_.

## 4 Simulation of Tug-of-War

In simulations of axonal transport, we record three states of the vesicle: pause, forward and backward (see Fig. 4) as follows: The *pause state* refers to situations when the cargo does not move or spends a short time (*<* 0.5 sec) in any direction before changing direction. The cargo is in *forward state* when it moves in an anterograde direction and in *backward state* when it moves in a retrograde direction for a longer period of time (*>* 0.5 sec).

**Figure 4.**
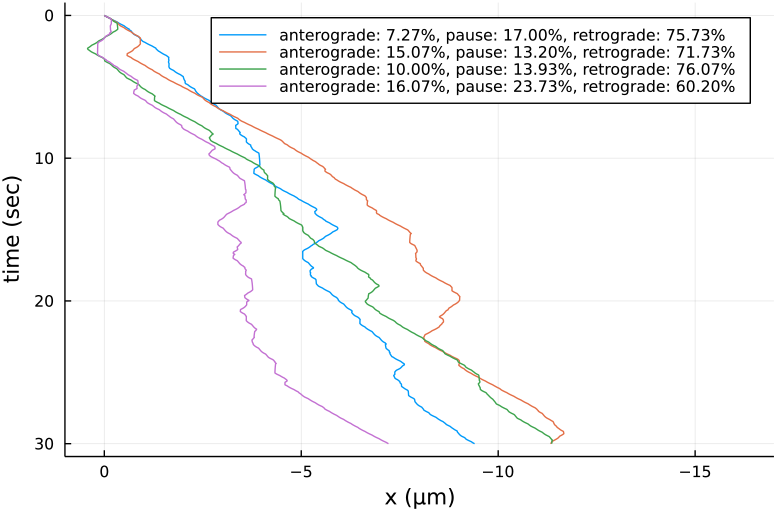
Simulated axonal transport for the reference parameters set (Table 1).

Fig. 4 shows that the cargo spends more time moving in retrograde direction than in pause or anterograde movement. Consequently, we refer to it as retrograde axonal transport that exhibits back-and-forth motion.

We run the simulation varying the parameters defined in the reference parameter set listed in Table 1. For the simulation results shown in Fig. 5 we vary the elasticity of actomyosin rings *κ* and their viscosity *χ* which gives rise to four different scenarios which we call “large *κ*”, “small *κ*”, “large *χ*’ and “small *χ*”.

**Figure 5.**
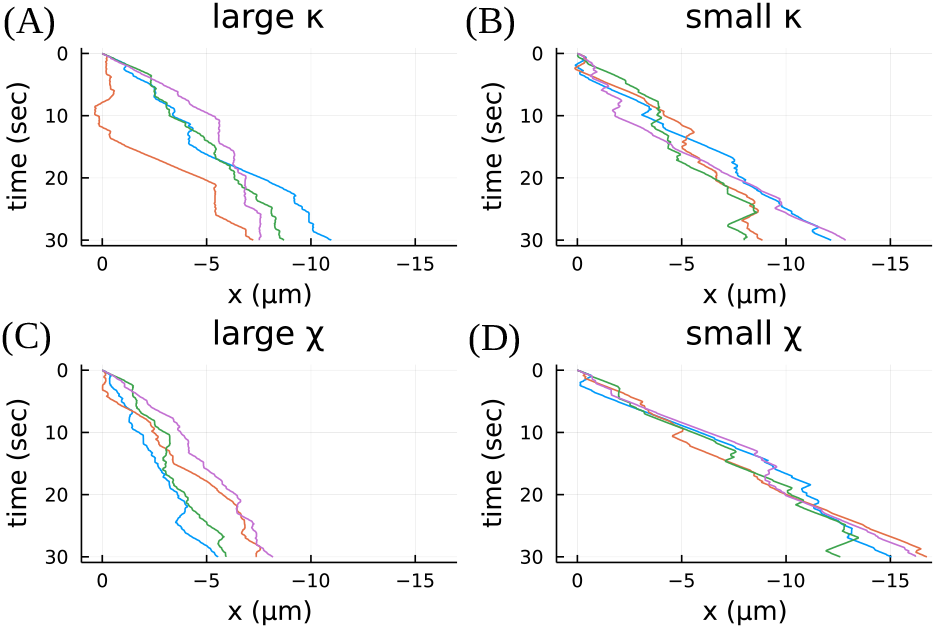
Simulations with modified parameter values for the elastic coefficient *κ* and the viscous coefficient *χ* of actomyosin rings. A: “large *κ*”: *κ →* 6 *× κ*, B: “small *κ*”: *κ →* 0.2 *× κ*, C: “large *χ*”: *χ →* 2.5 *× χ* and D: “small *χ*”: *χ →* 0.2 *× χ*.

We proceed similarly for variations of the reference values for axon radius *R*_*a*_ and drag friction *ξ* and show the simulations for the scenarios “large axon”, “small axon”, ‘large *ξ*” and “small *ξ*” in Fig. 6.

**Figure 6.**
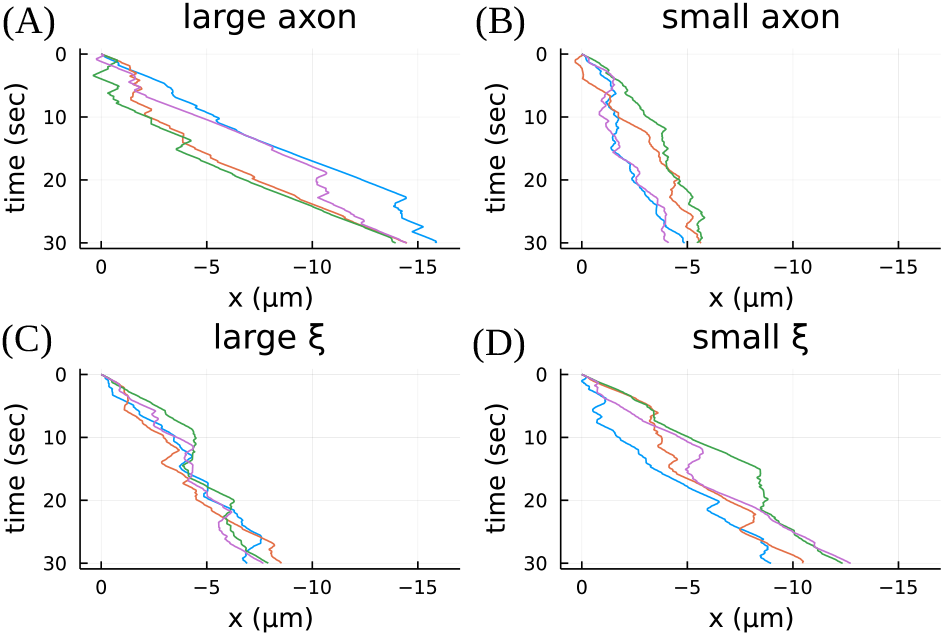
Simulations with modified parameter values for the axon radius *R*_*a*_ and the drag friction coefficient *ξ* of the vesicle. A: “large axon”: *R*_*a*_ *→* 1.25 *× R*_*a*_, B: “small axon”: *R*_*a*_ *→* 0.75 *× R*_*a*_, C: “large *ξ*”: *ξ →* 10 *× ξ* and D: “small *ξ*”: *ξ →* 0.1 *× ξ*.

For every parameter set discussed in Fig. 5 and Fig. 6, including the reference parameter set, we run *n* = 30 simulations visualising the distributions of the fractions of time spent in anterograde motion, pause and retrograde motion in Fig. 7 through boxplots (vertical lines: median and upper/lower quartile). We also list the average values in Table 2. In particular, we note concerning anterograde motion (Fig. 7A) that the elastic coefficient of actomyosin rings *κ* has the biggest impact with larger values increasing the time spent in anterograde motion. This indicates that the rigidity of actomyosin rings destabilises the dominant retrograde motion. This is also indicated by the fact the higher values of *κ* also increase the fraction of time spent in pause ((Fig. 7B) and significantly reduce the time spent in retrograde motion ((Fig. 7C).

**Figure 7.**
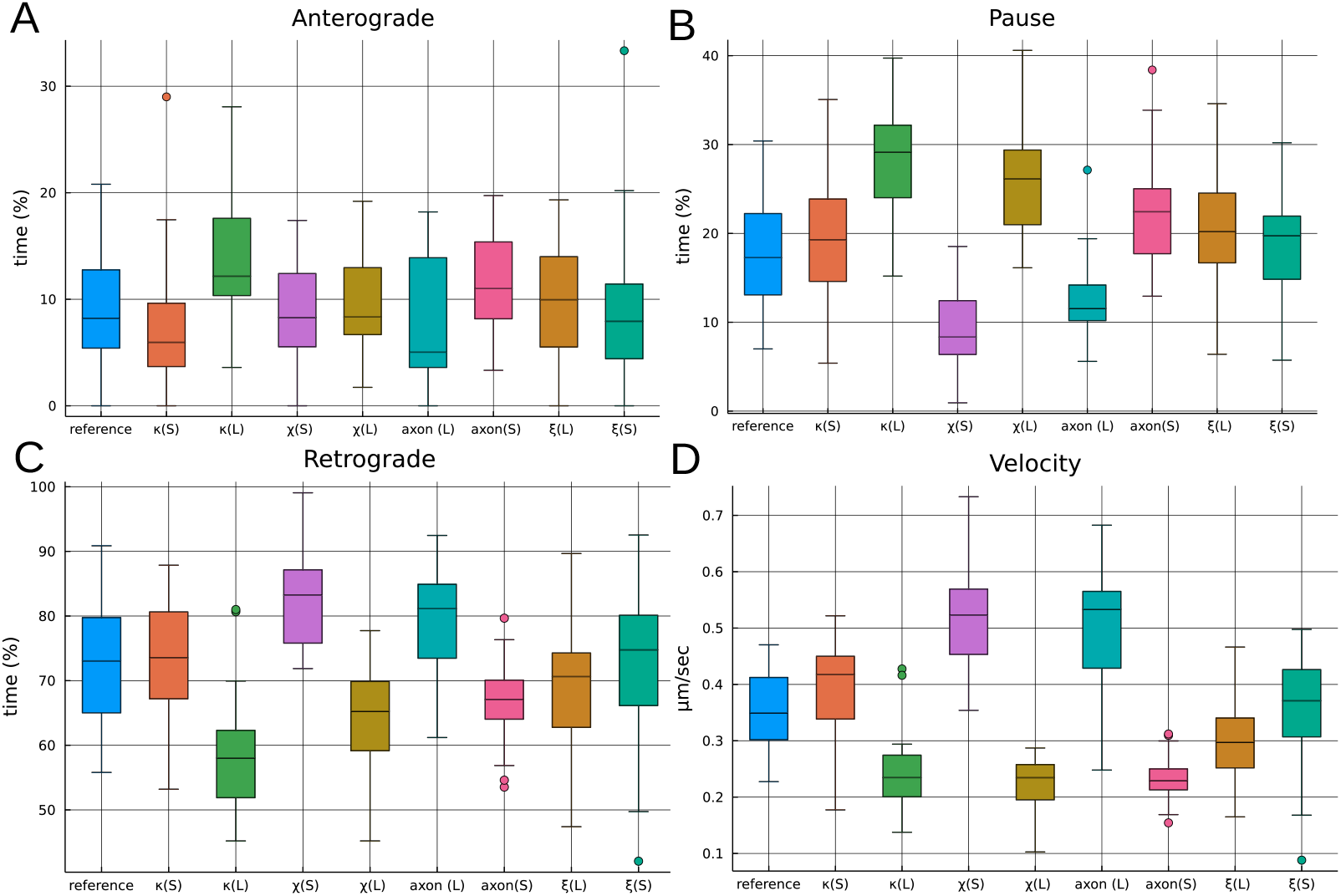
Comparison of reference parameter set and variations of actomyosin ring elasticity *κ* (“small *κ*(S)”, “large *κ*(L)”), actomyosin ring viscosity *χ* (“small *χ*(S)”, “large *χ*(L)”), actomyosin ring radius *R*_*a*_ (“small axon (S)”, “large axon (L)”) and vesicle drag friction *ξ* (“small *ξ*(S)”, “large *ξ*(L)”). A: Fraction of time spent in anterograde motion. B: Fraction of time spent in pause. C: Fraction of time spent in retrograde motion. D: Average velocities taken over the entire simulation run.

Concerning the fraction of time spent in pause (Fig. 7B) we note that also the viscosity *χ* of actomyosin rings has a significant impact. Larger values increase the fraction of time spent in pause at the expense of the fraction of time spent in retrograde motion (Fig. 7C). There is no significant impact on the time spent in anterograde motion, though, indicating that actomyosin ring viscosity may destabilize retrograde motion, but not stabilise anterograde motion.

We observe a similar response when varying the diameter of the axon. Larger axons significantly reduce the time spent in pause (Fig. 7B) mostly at the expense of the fraction of time spent in the dominant retrograde motion (Fig. 7C).

We now address the primary goal of this study which is to explain the observations about the effect of blebbistatin (BLB) on axonal transport reported in [15] and listed in the introduction. In the context of our model, we may model the effect of BLB in several different ways. In [15] it has been indicated that BLB increases the diameter of axons, therefore modelling BLB through increased values of *R*_*a*_ in our model is one possibility to mimic the effect of BLB. Alternatively, we may also assume that reduced force generation and cross-linking through myosin upon application of BLB should affect the mechanical properties of actomyosin rings and lead to either a reduced coefficient of elasticity *κ* and/or a reduced viscosity *χ* of actomyosin rings. This implies that the scenarios which should correlate with the application of BLB are axon(L), *κ*(S) and *χ*(S).

Our simulations (Fig. 7 and Table 2) indicate that for larger values of the axon radius *R*_*a*_ our model predicts indeed a decrease in the overall time spent in pause and an increase in the overall velocity of the vesicle – thus coinciding with three of the four observations we list in the introduction. The same effect can be observed for reduced viscosity *χ* and even for reduced *κ* in comparison with *κ*(L). Therefore, we argue that our model explains both observations.

Our model, however, so far fails at explaining that upon application of BLB, an increased fraction of time is spent in anterograde motion as compared to control. According to (Fig. 7) and Table 2) this fraction is predicted to be slightly smaller in all the scenarios which might correlate with BLB, i.e. for enlarged axons (axon(L)) and also for smaller values of either viscosity *χ* or elasticity coefficient *κ*. As a consequence, we cannot explain this effect based on the model we have formulated so far. In what follows, it is our goal to extend our model for axonal cargo transport by a minimal model for axon-cargo interaction in a way such that the modified model correctly predicts the results reported in [15].

## 5 Modelling contractile feedback of actin ring stretching

In this section we seek to introduce a minimal extension of the model introduced above which is sufficient to explain the increased fraction of time spent in anterograde motion upon inhibition of myosin motor proteins through blebbistatin (BLB). The model introduced above predicts a reduction of the time spent in pause upon myosin inhibition compensated by a larger fraction of time spent in the dominant retrograde direction. As for anterograde motion, the sensitivity analysis Fig. 7 reveals a rather negative trend upon myosin inhibition

This indicates that in order to correctly predict also increased anterograde motion upon myosin inhibition, our model lacks a mechanism that stabilises the direction of vesicles which occasionally might turn into anterograde direction. We argue that such stabilisation could originate from mechanotransductive positive feedback between the stress in the transversal actomyosin rings and myosin contraction. The stabilisation would be a consequence of myosin activation in stretched actomyosin rings which has been observed in other contexts [25]. Through such reinforcement, myosin contraction would be particularly effective in those actomyosin rings which are at the trailing end of the vesicle. Their contraction would push the vesicle forward and thus prevent the vesicle from pausing and eventually turning.

To formulate this as a system of equations, we assume that actomyosin rings undergo mechanotransduction reinforcing their stiffness temporarily upon stretching through myosin activation [25]. We introduce the myosin strength *m*_*j*_ of ring *j* and assume that it is upregulated upon stretching ring *j* and that in the absence of stretching it reverts to its base value with rate *β*,

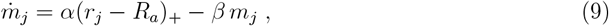

where *β* is the decay rate of myosin strength. We further assume that the coefficient of elasticity is given by a linear function of *m*_*j*_ which we write as

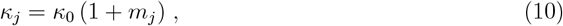

with the base elasticity coefficient *κ*_0_. Simulations of a steadily moving vesicle in response to a given, fixed pulling force replacing the pulling force exerted by molecular motors confirm that the myosin strength *m*_*j*_ is typically small away from the cargo and maximal at the trailing end of the cargo which we confirmed in simulations (Fig 8).

**Figure 8.**
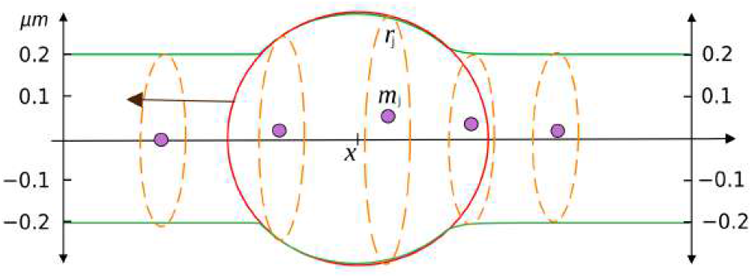
The scattered points represent the myosin strength *m*_*j*_ of the rings with radius *r*_*j*_ in a vesicle moving steadily in anterograde (left) direction.

We should also expect, that the force required to stretch the transverse actomyosin rings at the front end of the cargo vesicle opposes its motion. Yet, we expect this effect to be stronger in anterograde direction than in the dominant retrograde direction. For that reason, we model myosin inhibition through blebbistatin through a reduced base value for actomyosin ring elasticity, which we denote by *κ*^*−*^. Simulations of cargo trajectories in this scenario(Fig. 9B) with parameter values listed in Table 3 confirm that stretching induced myosin stabilisation leads to longer sections spent in anterograde direction. We use a larger value denoted by *κ*^+^ for actomyosin ring elasticity in control. Simulations (Fig. 9A) confirm that this will mostly prevent cargo from moving in anterograde direction, but not in retrograde direction.

**Table 3:** List of parameters values for stretch actomyosin ring contraction. The details of their derivation are summarised in section 6.

| Description | Symbol | Value |
| --- | --- | --- |
| Stretching stiffness of actin rings (control) | $\kappa_0^+$ | 6 pN $\mu\text{m}^{-1}$ |
| Stretching stiffness of actin rings (BLB) | $\kappa_0^-$ | 0.6 pN $\mu\text{m}^{-1}$ |
| Growth rate of myosin strength | $\alpha$ | 40 $\text{sec}^{-1} \mu\text{m}^{-1}$ |
| Decay rate of myosin strength | $\beta$ | 3 $\text{sec}^{-1}$ |

**Figure 9.**
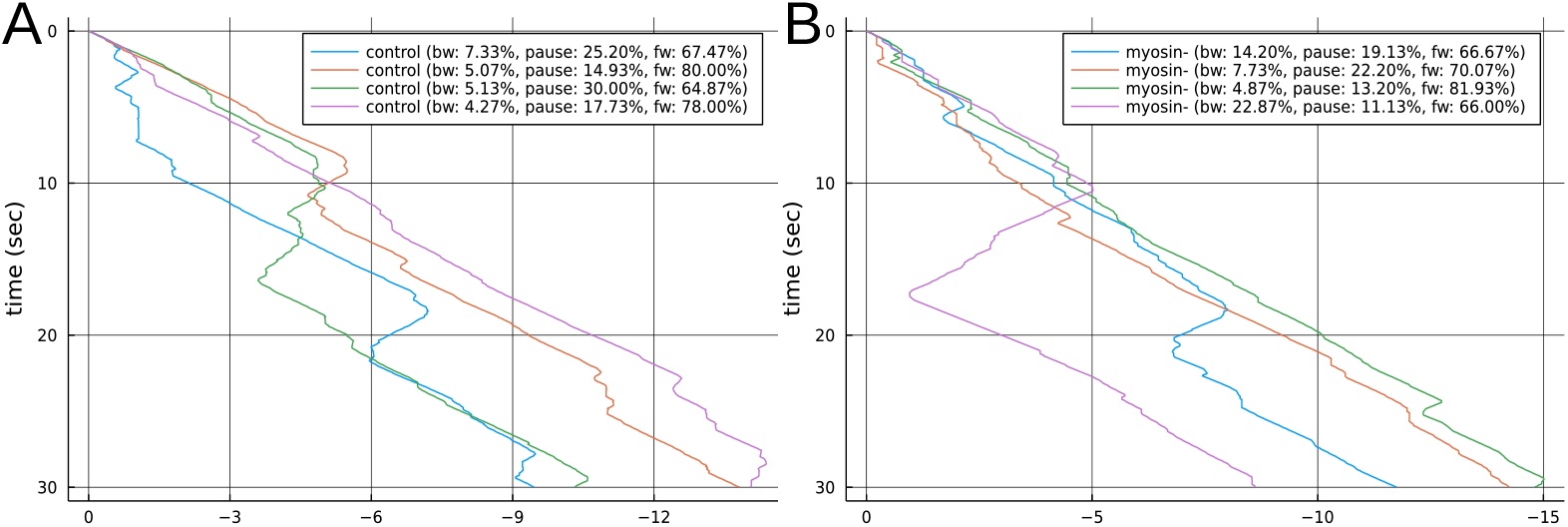
Simulations of axonal transport in control (A) and BLB myosin inhibition (B).

To quantify this effect we run a total of *n* = 30 simulations in both scenarios (Fig. 10). They show that with myosin inhibition the cargo vesicle spends a larger fraction of time (mean: 15.56%) moving in anterograde direction than in control (mean: 8.35%), which qualitatively coincides with the results reported in [15]. Pauses occur frequently in control (18.52% on average) compared to BLB myosin inhibition (mean: 11.64%). Finally, with myosin inhibition, the overall axonal speed increases on average by 0.40 µm sec^*−*1^. All this agrees well with the observations reported in [15] listed in the introduction.

**Figure 10.**
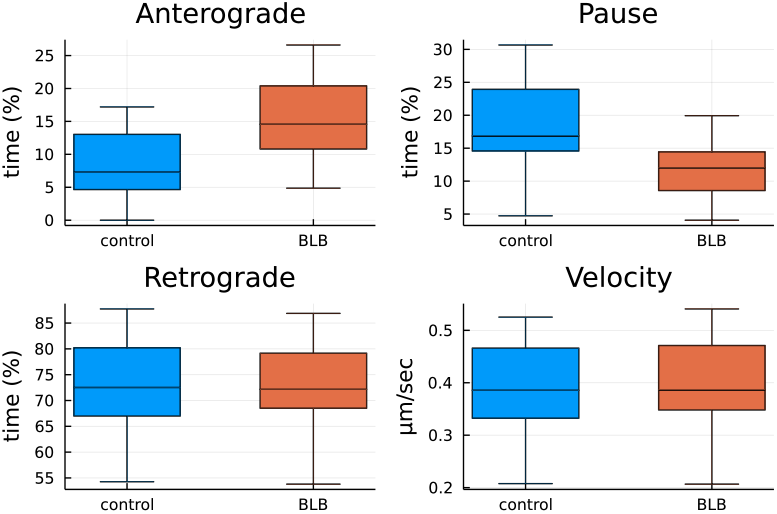
Comparison of simulations with transverse rings undergoing stretch-induced actomyosin contraction according to the parameter values listed in Table 3.

## 6 Model parameters

In this section, we discuss how we estimate those parameter values which are not reflected in the literature (see Table 1). For the coefficient for drag friction *ξ* we choose a relatively small value assuming that most of the drag will be exerted by the axonal cortex and the transverse actomyosin rings. Their effect is incorporated into our model through the separate parameters *κ* and *χ* for their elastic coefficient and viscous drag. Both values are chosen in a way such that the resulting forces acting on the cargo vesicle are in the 10 pN range which corresponds to the magnitude of pulling forces exerted by molecular motors.

As for the parameter values for the sub-model for stretch-induced myosin activation and elastic contraction (Table 3) which we describe in section 5, we determine the decay rate *β* in a way such that the characteristic timescale for myosin decay 1*/β* is a fraction of the time required to move past the distance *R*_*a*_ at a steady cargo velocity of about 0.5 µm sec^*−*1^ (see figure 1A and [15]). Using this approach we are able to achieve that the peak values of *m*_*j*_ are assumed at the transverse rings overlapping with the trailing part of the vesicle (Fig. 8).

The growth rate *α* in response to stretching, on the other hand, is determined in a way such that at peak stretching where *r*_*j*_ = *R*_*v*_ the equilibrium value of *m*_*j*_ where 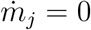 given by 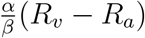 corresponds to the elastic coefficient *κ* in Table 1.

## 7 Conclusion

In this study it has been our goal to explain experimental observations on bidirectional transport of large cargo vesicles in axons in control and with myosin inhibition reported in [15]. We base our model on the mathematical framework for axonal cargo transport developed in [16] and incorporate the tug-of-war of motor proteins [7] pulling in anterograde and retrograde direction leading to bidirectional movement of cargo. The resulting model and simulations did predict some, but not all experimental observations. In particular, they would not predict a larger fraction of time spent in the non-dominant anterograde direction upon myosin inhibition.

We further incorporated mechanotransductive reinforcement [25] through stretch-induced up-regulation of contractile stress into the mechanical sub-model for transverse actomyosin rings. This acts as a minimal model stabilising the steady motion of cargo vesicles in either direction, but only when the overall level of actomyosin ring contraction is sufficiently low which is likely to be induced by myosin inhibition. In control, the model therefore predicts that cargo predominantly moves in retrograde direction in agreement with observations. In myosin inhibited cells, the model also correctly predicts the presence of periods spent in the non-dominant anterograde direction with the overall time spent in pause being reduced.

The reason is that in this sub-model transverse actomyosin rings have two roles. They exert elastic and viscous forces which oppose motion in any direction. In control cargo vesicles therefore predominantly undergo motion in the dominant retrograde direction. In cells in which myosin is inhibited, however, the contractile strength of the transverse rings increases through the stretching imposed by the large vesicle. During long-term steady motion the contractile strength peaks at the trailing end of the vesicle and the pushing forces the transverse rings exert prevent the vesicle from reverting into pause.

These results are consistent with the previous experimental results reported in [15]. We also confirm that the overall cargo velocity in BLB-treated cells is slightly elevated compared to the control. Our model based on stretching-induced reinforcement of transverse ring contractility therefore enables us to explain all the observations on the effect of blebbistatin induced mysoin inhibition on the axonal transport of large vesicles reported in [15].

This indicates that transversal actin rings act as stabilisers of directional transport via a myosin-dependent mechanotransductive feedback loop. This theoretical result could be tested experimentally by interrupting the mechanotransduction leading to potential myosin activation in response to stretching.

## Acknowledgments

NR was supported by an RTP scholarship funded by the Australian government. DO was supported by ARC Discovery Project DP180102956.

